# Cutting out the middle clam: lucinid endosymbiotic bacteria are also associated with seagrass roots worldwide

**DOI:** 10.1101/2020.05.06.080127

**Authors:** Belinda C. Martin, Jen A. Middleton, Matthew W. Fraser, Ian P.G. Marshall, Vincent V. Scholz, Hannes Schmidt

## Abstract

Seagrasses and lucinid bivalves inhabit highly reduced sediments with elevated sulphide concentrations. Lucinids house symbiotic bacteria (*Ca*. Thiodiazotropha) capable of oxidising sediment sulphide, and their presence in sediments has been proposed to promote seagrass growth by decreasing otherwise phytotoxic sulphide levels. However, vast and productive seagrass meadows are present in ecosystems where lucinids do not occur. Hence, we hypothesised that seagrasses themselves host sulphur-oxidising bacteria that could secure their survival when lucinids are absent. We analysed newly generated and publicly available 16S rRNA gene sequences from seagrass roots and sediments across 14 seagrass species and 10 countries and found that persistent and colonising seagrasses across the world harbour sulphur-oxidising *Ca*. Thiodiazotropha, regardless of the presence of lucinids. We used fluorescence *in situ* hybridisation to visually confirm the presence of *Ca*. Thiodiazotropha on roots of *Halophila ovalis*, a colonising seagrass species with wide geographical, water depth range, and sedimentary sulphide concentrations. We provide the first evidence that *Ca*. Thiodiazotropha are commonly present on seagrass roots, providing a mechanism for seagrasses to alleviate sulphide stress globally.

## Introduction

Seagrasses are marine flowering plants that cover an estimated global area of 300 000 – 600 000 km^2^ and are crucial to the health of shallow coastal ecosystems worldwide [1]. Seagrasses typically thrive in highly reduced sediments where sulphide concentrations can accumulate to phytotoxic levels (> 10 μmol L^-1^ [2]), which presents an ongoing enigma as to how they survive. Bivalves belonging to the family Lucinidae are often abundant in seagrass sediments. Lucinids, unlike most animals, thrive in sulphide-rich sediments as they house symbiotic chemoautotrophic bacteria (*Ca*. Thiodiazotropha) inside their gills which oxidise sulphides to provide energy for CO_2_ fixation. Because of their ability to remove sulphides from the sediment, lucinids have been proposed as a mechanism for sulphide detoxification in seagrass beds globally [3]. In this “tripartite” relationship, seagrasses benefit from reduced sulphide intrusion, whilst lucinids and their symbionts profit from enhanced sulphide production arising from seagrass organic matter accumulation as well as oxygen leaking from actively growing seagrass roots and rhizomes. This relationship has been recently tested in the field, where increased densities of the lucinid *Loripes orbiculatus* correlated with reduced sulphide intrusion in the seagrass *Zostera noltei* [4]. Whilst this study provides compelling evidence for sulphide detoxification for seagrass systems with high densities of lucinid bivalves, it is not adequate for explaining sulphide detoxification for systems lacking lucinids.

Lucinids acquire their endosymbionts in free-living stage from the environment, believed to be a result of chance encounters with potential symbionts during the juvenile phase [5, 6]. It follows that seagrass roots, which provide a chemical environment that bears a striking resemblance to lucinid gills (i.e. gradients of oxygen, organic matter, sulphide and carbon dioxide availability) [7, 8], would also provide a suitable niche to attract and possibly attain these bacteria. As part of recolonisation of the marine environment, seagrasses require adaptions to cope with high sulphide environments, one of which could be supported via their root microbiota. Thus, we hypothesized that seagrass roots harbour sulphur-oxidising *Ca*. Thiodiazotropha, which could explain their successful colonisation even in sulphide-rich sediments where lucinids are absent.

## Methods

To explore the possibility that seagrass roots serve as a suitable habitat for *Ca*. Thiodiazotropha, we analysed 16S rRNA gene sequences (Illumina MiSeq platform) recovered from publicly available and newly generated data on seagrass roots and sediments across 14 seagrass species and 10 countries (Fig. 1). All raw sequences were processed through the DADA2 pipeline [9] using SILVA 132 database [10] to assign taxonomy. Amplicon Sequence Variants (ASVs) classified as *Ca*. Thiodiazotropha were compared to SILVA 138 using blastn (v 2.2.29+) [10] and a phylogenetic tree was constructed using PhyML [11] with 1000 bootstraps. We also applied fluorescence *in situ* hybridisation (FISH) to visualise *Ca*. Thiodiazotropha on roots of the seagrass *Halophila ovalis* following the protocol outlined in [8, 12] (detailed methods in supplementary).

**Figure 1.**
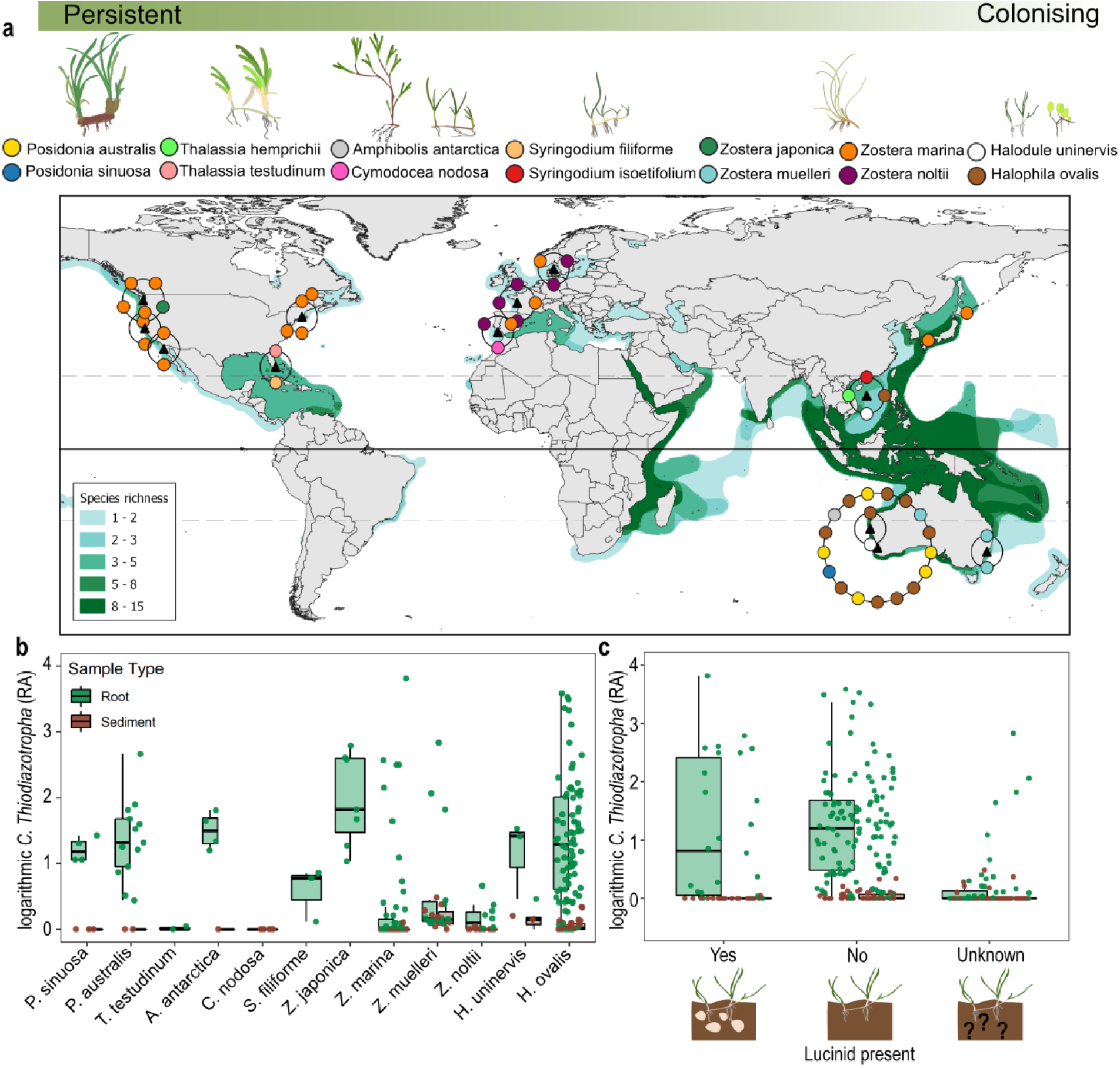
Global distribution of seagrass samples and the relative abundance of *C*. Thiodiazotropha. **(a)** Seagrass species and sample locations analysed in this study and the global distribution of seagrass species. Seagrasses are classified by functional type; with seagrasses on the left classified as persistent (large, slow growing and forms persistent meadows), and those on the right as colonising (small, fast growing and forms transient meadows). Global seagrass distribution data was acquired from Green and Short [14]. Black triangles indicate a cluster of sites and/or seagrass species that were collected in close proximity. Location meta-data can be found in Table S1 and Table S2. **(b)** Relative abundance of *Ca*. Thiodiazotropha sequences (logarithmic scale) recovered from all seagrass roots and sediments. **(c)** Relative abundance of *Ca*. Thiodiazotropha sequences (logarithmic scale) recovered from seagrass roots and sediments from environments with and without lucinids present (lucinid precense/absence references available in Table S2).

## Results and Discussion

We recovered *Ca*. Thiodiazotropha sequences from roots of 12 of the 14 seagrass species analysed, comprising four seagrass families and three independent lineages of seagrass evolution [13]. When present, *Ca. Thiodiazotropha* sequences were far more abundant in seagrass root samples compared to the surrounding sediment, where often no sequences were recovered at all (Fig. 1b). *Ca*. Thiodiazotropha sequences were also found in seagrasses of all functional types; from large, slow growing seagrass species that develop persistent meadows such as *Posidonia* spp. to smaller fast growing species that form transient meadows such as *Halophila* spp. (Fig. 1). Data are skewed towards temperate systems, however, *Ca*.

Thiodiazotropha sequences were also obtained from tropical seagrass roots in Florida, as well as the tropical Paracel Islands in the South China Sea. Notably, comparable levels of relative abundance of *Ca*. Thiodiazotropha from seagrass roots appeared independent of the presence or absence of lucinids (Fig. 1c). Together, this suggests that *Ca*. Thiodiazotropha colonise seagrasses irrespective of their geography, lifestyle, evolutionary history and proximity to lucinids likely because the root environment and its mosaic of chemical gradients (e.g. oxygen, pH, metals and nutrients) [7, 8] provides a suitable niche for these chemoautotrophs.

Most seagrass species exhibited a relatively diverse set of ASVs aligning to known endosymbionts of lucinid species (Fig. S1). Some lucinid species also exhibit diversity in their endosymbionts, possibly as a reflection of the diversity of free-living symbionts present in the environment [5, 6]. To date, functional analysis of three gill edosymbionts (including *Ca*. Thiodiazotropha endolucinida indicated in Fig. S1), revealed not only genes relating to chemoautotrophy, but also diazotrophy, heterotrophy and oxidation of C1 compounds [5, 15–17]. Hence these bacteria may not only provide a means for lowering sediment sulphide levels, but may also provide an additional source of NH_4_ ^+^ via nitrogen fixation; the preferred source of N for seagrass uptake [18].

FISH with a *Ca*. Thiodiazotropha-targeted oligonucleotide probe combined with class- and domain-level probes showed that *Ca*. Thiodiazotropha forms dense colonies on the surface of *Halophila ovalis* roots, particularly in axial grooves between epidermal cells and on the base of root hairs (Fig 2, Fig S2). Strikingly, a morphological diversity of *Ca*. Thiodiazotropha was apparent in FISH images – with smaller single-cell rods (∼1µm) present on the root surface (e.g. Fig S3a), as well as larger coccoid shaped (∼2 µm) cells (e.g. Fig S3d) that form dense colonies in axial grooves and root hairs. Such differences in morphology have been previously observed between extracellular (those found in the sediment) and intracellular lifestyles (those located inside lucinid gills) for the symbionts of the lucinid *Codakia orbicularis* [19]. We cannot say if our observations of *Ca*. Thiodiazotropha is a reflection of morphological plasticity or closely related strains with differing morphologies. Regardless, we provide the first *in situ* evidence that *Ca*. Thiodiazotropha is colonising seagrass roots in high abundances. Together with amplicon sequencing based evidence that *Ca*.

Thiodiazotropha are associated with seagrasses of varying life-strategies and evolutionary histories worldwide, our data suggests that this relationship may be both general and essential for seagrasses to thrive in sulphide-rich sediments, thus extending the proposed lucind-seagrass relationship.

**Figure 2:**
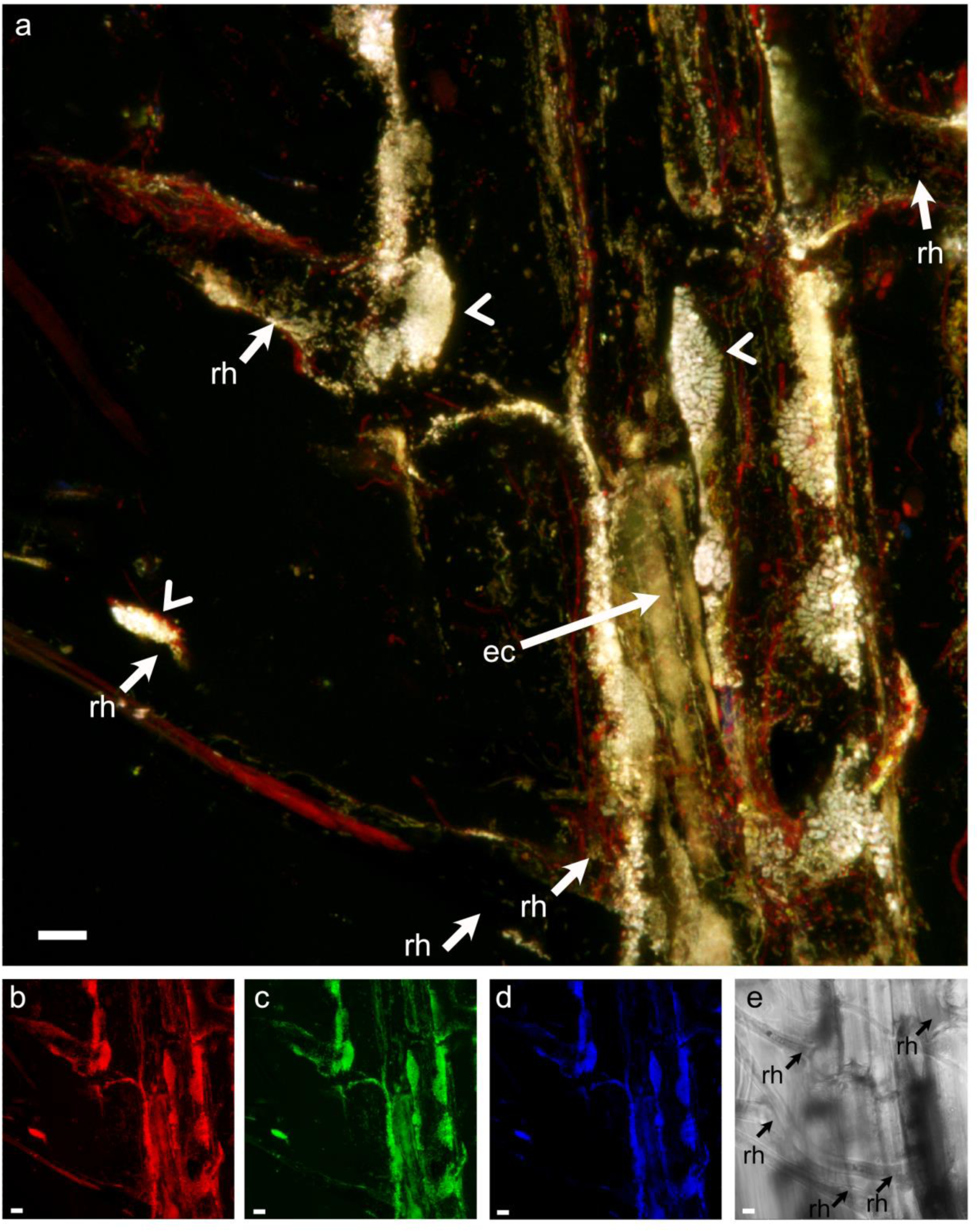
Projected images of *Halophila ovalis* roots (collected from the Swan River, Western Australia) with associated populations of *Ca*. Thiodiazotropha. **a** overlay of all probes. Large clusters of *Ca*. Thiodiazotropha can be seen in yellow/white (white arrow heads) and populations of other bacteria (particularly filamentous bacteria) can be seen in red. Several root hairs are present (rh; white arrow) and the epidermal cells of the seagrass are also visible (ec; white arrow). N.B. *H. ovalis* possesses no lateral roots, so all ‘branching’ are root hairs (i.e. not lateral roots). **b** *Bacteria* (probe: EUBI-III-ATTO 647), **c** *Gammaproteobacteria* (probe: GAM42a-ATTO 565), **d** *Ca*. Thiodiazotropha (probe: THIO-847) **e** transmission image with the bases of root hairs (rh) indiacted with black arrows. Scale bars: 10 µm.

## Acknowledgements

The authors wish to thank Gary Kendrick, Jeremy Bougoure, Daniela Trojan, PWIS and PP for advice and fruitful discussions. MWF was supported by the Robson and Robertson postdoctoral fellowship awarded by the UWA Oceans Institute. This research was partly supported by a Collaborative Research Grant awarded by the Indian Ocean Marine Resaerch Centre, and Strategic Funding awarded by the School of Biological Sciences, UWA. The authors acknowledge the facilities, and the scientific and technical assistance of Microscopy Australia at the Centre for Microscopy, Characterisation & Analysis, The University of Western Australia, a facility funded by the University, State and Commonwealth Governments.

## Conflict of interest

All authors declare there are no competing financial interests in relation to the work described.

## Supplementary

**Figure S1.**
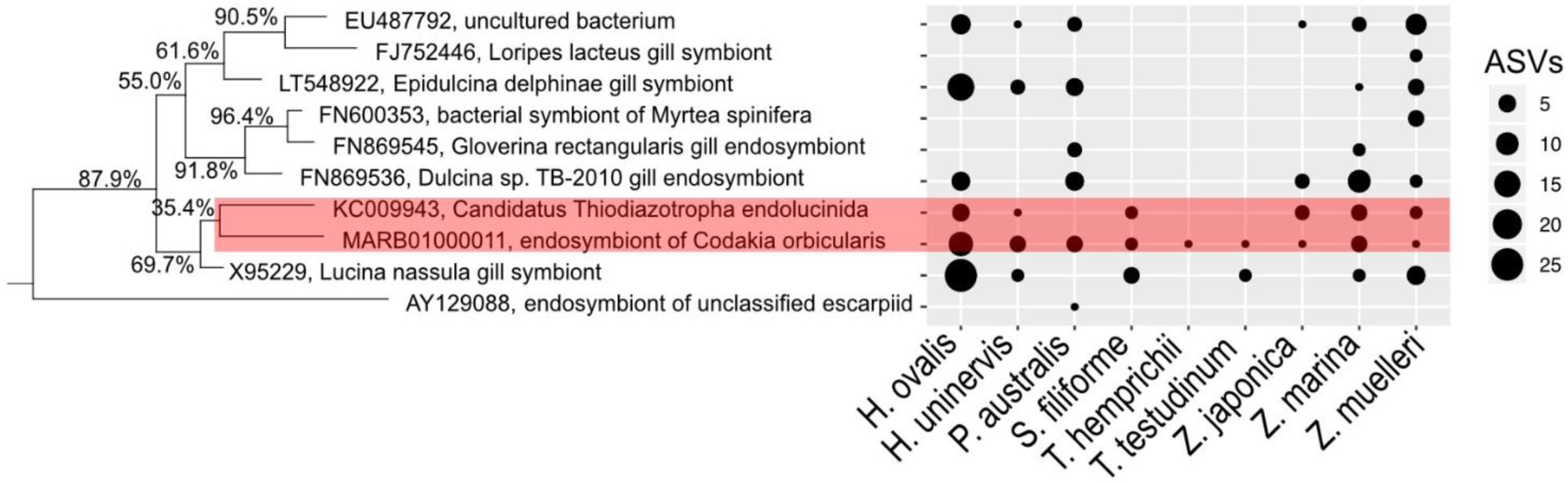
BLAST best hits to the SILVA 138 database of the 180 *Ca*. Thiodiazotropha ASVs identified in this study, organised according to a phylogenetic tree constructed from the full-length 16S rRNA gene sequences indicated. 175/180 (97.2%) of all ASVs were >97.0% identical to the full-length reference sequence indicated in the phylogenetic tree. The remaining 5 ASVs were all >93.9% identical to the full-length reference sequences. Bootstrap percentages are indicated on the phylogenetic tree.

**Figure S2.**
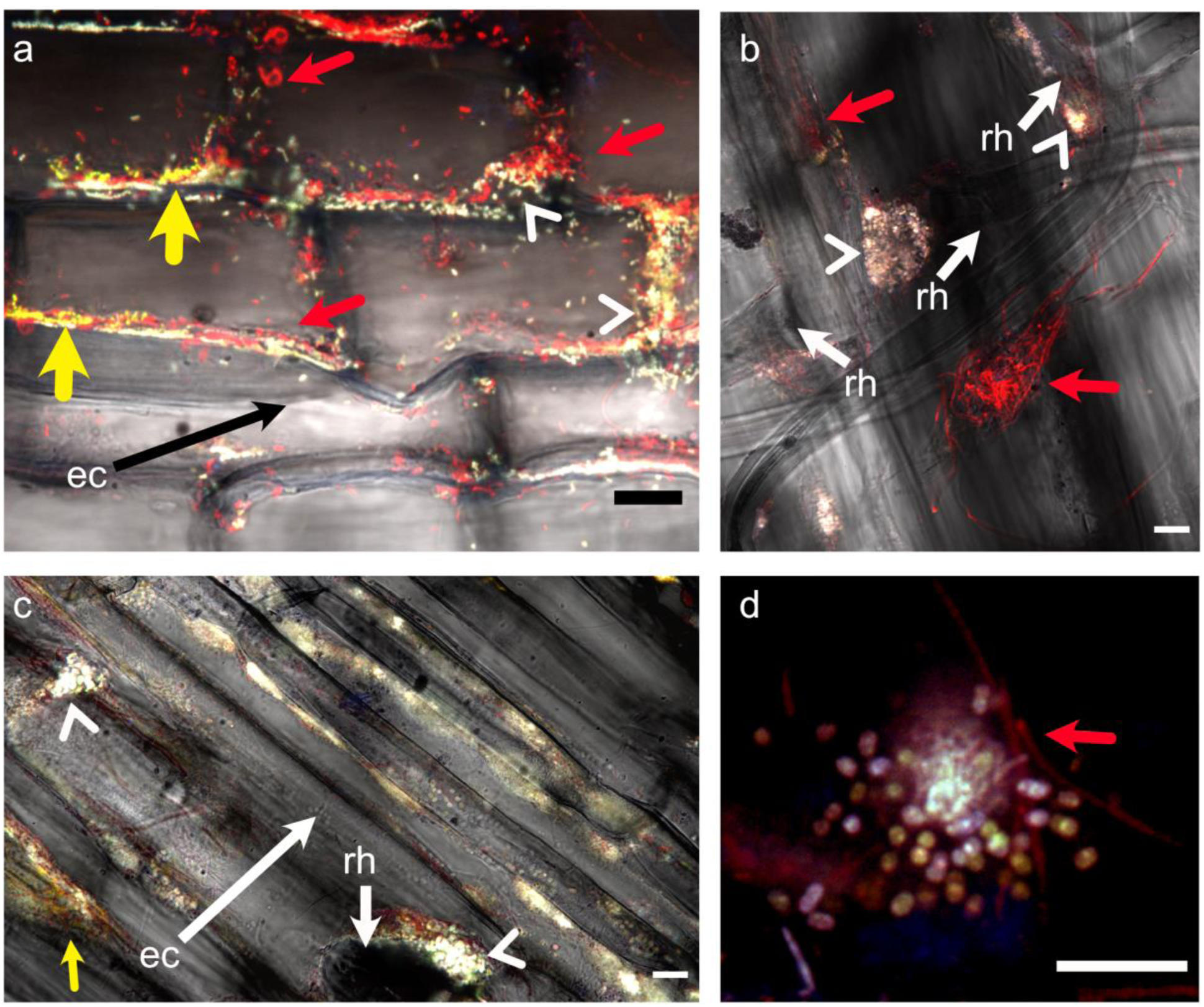
Representative confocal laser scanning microscopy (CLSM) micrographs of *Halophila ovalis* roots hybridised with bacteria (EUBI-III), *Gammaproteobacteria* (GAM42a) and *Ca. Thiodiazotropha* (THIO-845). **a** overlay of all probes and transmission image showing populations of *Ca. Thiodiazotropha* (white arrow heads) within the axial groves between epidermal cells (ec). Other populations of *Gammaproteobacteria* can be seen in yellow (yellow arrows) and other bacteria in red (red arrows). **b** overlay of all probes and transmission image showing populations of *Ca. Thiodiazotropha* (white arrow heads) clustering at the base of root hairs (rh). Other clusters of bacteria can be seen in red (red arrows). **c** overlay of all probes and transmission image showing populations of *Ca. Thiodiazotropha* (white arrow heads) within the axial groves between epidermal cells (ec) and bases of root hairs (rh). **d** Magnified image of coccoid shaped *Ca. Thiodiazotropha* (yellow/pink). Other filamentous bacteria can be seen in red. Scale bars: 10 µm.

## Extended methods

### 16S rRNA gene data analysis

Primers were removed from all raw 16S rRNA gene data using cudadapt (v. 2.7) [20]. The trimmed fastq files were then analyzed in R (v. 3.6.1) using dada2 (v. 1.30.0), phyloseq (v. 1.30.0) and ggplot2 (v. 3.1.0) [9, 21, 22]. Taxonomy was assigned using the SILVA 132 short subunit database [10]. Sequences assigned as “Chloroplast” and “Mitochondria” were removed from each individual study. Sequences assigned as “*Candidatus* Thiodiazotropha” from each study were then calculated as both total reads and as relative abundance of total reads. Control samples (where available) were also checked for the occurrence of “*Candidatus* Thiodiazotropha”. Amplicon Sequence Variants (ASVs) classified as *Ca*. Thiodiazotropha were compared to SILVA 138 NR 99 short subunit database using blastn version 2.2.29+ [10]. Top hit sequences, pre-aligned in SILVA, were used to make a phylogenetic tree using PhyML with 1000 bootstraps [11].

### FISH-CLSM

Standard FISH was conducted on four replicate unwashed root segments (∼ 1 cm, of the seagrass *Halophila ovalis* that had been collected from the Swan River, Western Australia (−32.03, 115.76) and fixed in 4% formaldehyde (v/v) over night at 4 °C. A subset of these four replicate root samples were also sequenced (16S rRNA gene, Illumina MiSeq platform; study PRJNA545746 in Table S1).

Probe specificity of the “*Ca*. Thiodiazotropha” probe THIO-845 (Table S1) was checked using TestProbe [10] against the SILVA 138 NR with zero mismatches. The abundance of reads from outgroup matches, as well as the target *Ca*. Thiodiazotropha were then summed across the four replicate samples that had been sequenced to check for potential off-target hybridisations (Table S2). The bacterial probe mix (EUB338I-III) combined with the *Gammaproteobacteria* probe (GAM42a) were used as positive controls, whilst the NON338 probe (one for each fluorophore) served as a negative control. An additional negative control was performed on the biofilm of fixed *Eucalyptus camadulensis* leaves (with no *Ca*. Thiodiazotropha present as indicated by 16S rRNA gene sequencing) using the THIO-845 probe in combination with EUBmix and GAM42a to check for off-target hybridisations.

For the hybridisations, roots were placed in 1mL tubes and incubated in 540 µL of hybridisation buffer (900 mM NaCl, 20 mM Tris-HCl, 35% formamide (v/v), 10% SDS (v/v)) and 15 µL of each probe (50 ng µL^-1^)[12]. All probes were synthesised by biomers.net (Ulm, Germany) and were hybridised at a formamide concentration of 35% at 46 °C for 3.5 h (Table S3). Root sections were carefully transferred to 50 mL of pre-warmed washing buffer (70 mM NaCl, 20 mM Tris-HCl, and 5 mM EDTA) and incubated for 10 min at 48 °C. Root pieces were then transferred to cold MQ H_2_O for 1 min before mounting in VectaShield AF1 (Vector Laboratories, Burlingame, CA) on a glass slide.

**Table S1.**
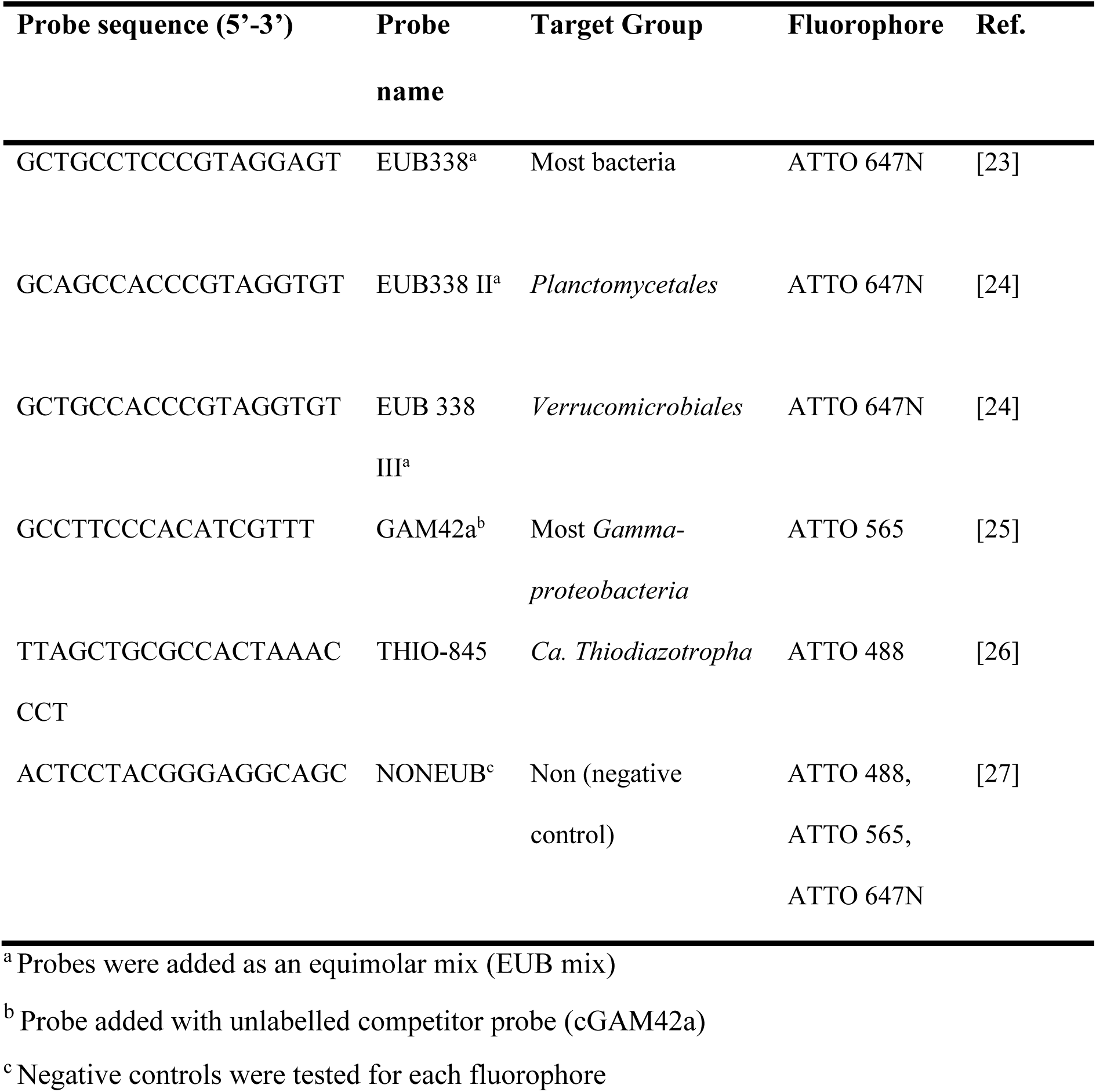
Fluorescently labelled oligonucleotide probes applied in this study.

**Table S2.** Top ten taxa from ProbeTest result for THIO-845 and number of sequences recovered from 16S rRNA gene sequencing

| Taxa | Coverage | Specificity | Acc. | Eligible | Match | Mismatch | # seqs |
| --- | --- | --- | --- | --- | --- | --- | --- |
| Chromatiaceae;<br>Phaeochromatium | 100 | 100 | 8 | 8 | 8 | 0 | 0 |
| **Sedimenticolaceae; Ca**<br>**Thiodiazotropha** | **96.2** | **100** | **211** | **211** | **203** | **8** | **52,331** |
| Chromatiaceae;<br>Marichromatium | 96.1 | 100 | 77 | 77 | 74 | 3 | 0 |
| Chromatiaceae;<br>Halochromatium | 92.1 | 100 | 354 | 354 | 326 | 28 | 0 |
| Gammaproteobacteria;<br>SZB30– | 91.7 | 100 | 354 | 354 | 326 | 28 | 0 |
| Nitrococcales;<br>Nitrococcaceae | 87.9 | 100 | 33 | 33 | 29 | 4 | 0 |
| **Chromatiaceae; uncultured** | **59.9** | **100** | **42** | **42** | **25** | **17** | **1,747** |
| **Arenicellaceae; Ca**<br>**Thiozymbion** | **55.6** | **100** | **108** | **108** | **60** | **48** | **140** |
| Gammaproteobacteria;<br>TDNP-Wbc97-11-7-16 | 50 | 100 | 2 | 2 | 1 | 1 | 0 |
| **Gammaproteobacteria;**<br>**Endothiovibrio** | **33.3** | **100** | **3** | **3** | **1** | **2** | **49** |

Confocal laser scanning microscopy (CLSM) was performed on a Nikon Ti-E inverted microscope with a Nikon A1Si spectral detector. Fluorescent dyes Atto 488, Atto 565, and Atto 647N were sequentially excited with 488, 561, and 640 nm laser lines, respectively. Laser power was set at 4% and photomultiplier gain and offset were individually optimised for each channel and every field of view. For each sample, the upper root surface was located and images were acquired using a × 60 Plan Apo VC Oil lens with four-line averaging. Image stacks (z = 0.6 µm) were sequentially acquired for all laser lines and projected into single layers and three-dimensional images using NIS Elements Viewer © (version 5.21.00).

## Competing interests

There are no competing financial interests in relation to the work described

## Notes

### Competing Interest Statement

The authors have declared no competing interest.

## References

1. Orth RJ, Carruthers TJB, Dennison WC, Duarte CM, Fourqurean JW, Heck KL, et al. A global crisis for seagrass ecosystems. Bioscience 2006; 56: 987–996.

2. Lamers LPM, Govers LL, Janssen ICJM, Geurts JJM, Van der Welle MEW, Van Katwijk MM, et al. Sulfide as a soil phytotoxin-a review. Front Plant Sci 2013; 4: 268.

3. Van der Heide T, Govers LL, de Fouw J, Olff H, Van der Geest M, Van Katwijk MM, et al. A three-stage symbiosis forms the foundation of seagrass ecosystems. Science (80-) 2012; 336: 1432–1434.

4. Geest M Van Der, Heide T Van Der, Holmer M, Wit R De. First Field-Based Evidence That the Seagrass-Lucinid Mutualism Can Mitigate Sulfide Stress in Seagrasses. Front Mar Sci 2020; 7: 1–13.

5. Lim SJ, Alexander L, Engel AS, Paterson AT, Anderson LC, Campbell BJ. Extensive Thioautotrophic Gill Endosymbiont Diversity within a Single Ctena orbiculata (Bivalvia: Lucinidae) Population and Implications for Defining Host-Symbiont Specificity and Species Recognition. mSystems 2019; 4: 1–19.

6. Brissac T, Merçot H, Gros O. Lucinidae/sulfur-oxidizing bacteria: Ancestral heritage or opportunistic association? Further insights from the Bohol Sea (the Philippines). FEMS Microbiol Ecol 2011; 75: 63–76.

7. Brodersen KE, Koren K, Moßhammer M, Ralph PJ, Kühl M, Santner J. Seagrass-mediated phosphorus and iron solubilization in tropical sediments. Environ Sci Technol 2017; 51: 14155–14163.

8. Martin BC, Bougoure J, Ryan MH, Bennett WW, Colmer TD, Joyce NK, et al. Oxygen loss from seagrass roots coincides with colonisation of sulphide-oxidising cable bacteria and reduces sulphide stress. ISME J 2019; 13: 707–719.

9. Callahan BJ, McMurdie PJ, Rosen M, Han AW, Johnson AJA, Holmes S. DADA2: High resolution sample inference from Illumina amplicon data. Nat Methods 2016; 13: 4–5.

10. Quast C, Pruesse E, Yilmaz P, Gerken J, Schweer T, Yarza P, et al. The SILVA ribosomal RNA gene database project: Improved data processing and web-based tools. Nucleic Acids Res 2013; 41: 590–596.

11. Guindon S, Gascuel O. A Simple, Fast, and Accurate Algorithm to Estimate Large Phylogenies by Maximum Likelihood. Syst Biol 2003; 52: 696–704.

12. Schmidt H, Eickhorst T. Detection and quantification of native microbial populations on soil-grown rice roots by catalyzed reporter deposition-fluorescence in situ hybridization. FEMS Microbiol Ecol 2014; 87: 390–402.

13. Les DH, Cleland MA, Waycott M. Phylogenetic Studies in Alismatidae, II : Evolution of Marine Angiosperms (Seagrasses) and Hydrophily. Am Soc Plant Taxon 1997; 22: 443–463.

14. Green EP, Short FT. World atlas of seagrasses. 2003. Prepared by UNEP World Conservation Monitoring Centre, Berkeley, California, USA.

15. Petersen JM, Kemper A, Gruber-Vodicka H, Cardini U, Van Der Geest M, Kleiner M, et al. Chemosynthetic symbionts of marine invertebrate animals are capable of nitrogen fixation. Nat Microbiol 2016; 2: 1–11.

16. König S, Gros O, Heiden SE, Hinzke T, Thürmer A, Poehlein A, et al. Nitrogen fixation in a chemoautotrophic lucinid symbiosis. Nat Microbiol 2016; 2.

17. Lim SJ, Davis BG, Gill DE, Walton J, Nachman E, Engel AS, et al. Taxonomic and functional heterogeneity of the gill microbiome in a symbiotic coastal mangrove lucinid species. ISME J 2019; 13: 902–920.

18. Touchette BW, Burkholder JM. Review of nitrogen and phosphorus metabolism in seagrasses. J Exp Bot 2000; 250: 133–167.

19. Gros O, Liberge M, Heddi A, Khatchadourian C, Felbeck H. Detection of the Free-Living Forms of Sulfide-Oxidizing Gill Endosymbionts in the Lucinid Habitat (Thalassia testudinum Environment). Appl Environ Microbiol 2003; 69: 6264–6267.

20. Martin M. Cutadapt removes adapter sequences from high-throughput sequencing reads. EMBnetJ 2011; 7: 2803–2809.

21. McMurdie PJ, Holmes S. Phyloseq: an R package for reproducible interactive analysis and graphics of microbiome census data. PLoS One 2013; 8: 1–11.

22. Wickham. ggplot2: Elegant Graphics for Data Analysis. 2009. Berlin: Springer Science & Business Media.

23. Amann RI, Blinder BJ, Olson RJ, Chisholm SW, Devereux R, Stahl DA. Combination of 16S rRNA-targeted oligonucleotide probes with flow cytometry for analyzing mixed microbial populations. Appl Environ Microbiol 1990; 56: 1919–25.

24. Loy A, Lehner A, Lee N, Adamczyk J, Meier H, Ernst J, et al. Oligonucleotide microarray for 16S rRNA gene based detection of all recognized lineages of sulfate reducing prokaryotes in the environment. Appl Env Microbiol 2002; 68 SRC-: 5064–5081.

25. Alm EW, Oerther DB, Larsen N, Stahl DA, Raskin L. The oligonucleotide probe database. Appl Environ Microbiol 1996; 62: 3557–3559.

26. Hausl BR. Genome diversity and free-living lifestyle of chemoautotrophic lucinid symbionts. 2017. University of Vienna.

27. Wallner G, Amann R, Beisker W. Optimizing fluorescent in situ hybridization with rRNA targeted oligonucleotide probes for flow cytometric identification of microorganisms. Cytometry 1993; 14: 136–143.

